# Modulation of sweet preference by neurosteroid-sensitive, δ-GABA_A_ receptors in adult mouse gustatory insular cortex

**DOI:** 10.1101/2024.10.04.616671

**Authors:** Priscilla E Yevoo, Alfredo Fontanini, Arianna Maffei

## Abstract

Taste preference is a fundamental driver of feeding behavior influencing dietary choices and eating patterns. Extensive experimental evidence indicates that the gustatory cortex (GC) is engaged in taste perception, palatability and preference. However, our knowledge of the neural and neurochemical signals regulating taste preference is rather limited. Neuromodulators can affect preferences, though their effects on neural circuits for taste are incompletely understood. Neurosteroids are of particular interest in view of reports that systemic administration of the neurosteroid allopregnanolone, a potent modulator of tonic GABAergic inhibition, induces hyperphagia and increases intake of energy rich food in human and animal subjects. Tonic inhibition is a powerful modulator of circuit excitability and is primarily mediated by extrasynaptic GABA_A_ receptors containing the delta subunit (δ-GABA_A_Rs). These receptors are widely distributed in the brain, but information regarding the expression of δ-GABA_A_Rs within gustatory circuits is lacking, and their role in taste preference has not been investigated. Here, we focused on GC to investigate whether activation of δ-GABA_A_Rs affects sweet taste preference in adult mice. Our data reveal that δ-GABA_A_Rs are expressed in multiple cell types within GC. These receptors mediate an allopregnanolone-sensitive tonic current and decrease sweet taste preference by altering the behavioral sensitivity to sucrose concentration in a cell type-specific manner. Our findings demonstrate that taste sensitivity and preference in the adult mammalian brain are modulated by tonic inhibition mediated by neurosteroid-activated δ-GABA_A_Rs in GC.

## Introduction

Taste plays a crucial role in shaping feeding behaviors, with foods perceived as palatable consumed more than foods perceived as bland or unpleasant. Abnormal eating patterns such as hyperphagia or food avoidance are associated with altered taste perception and changes in taste preference. In addition, imaging studies of subjects with eating disorders or other conditions associated with abnormal eating habits reported altered activity in the gustatory insular cortex (GC)^1–3^, a region involved in taste processing^4,5^, palatability^6,7^, preference^8^ and taste-related behaviors^9–11^. These findings suggest a direct relationship between GC circuits and modulation of taste preference. Extensive work has explored the role of neuromodulators on reward circuits engaged in food choices^12,13^. However, neurochemical signals regulating taste preferences and the contribution of GC neurons in this process remain unclear.

Neurosteroids, allosteric modulators of neurotransmission produced in an activity-dependent manner^14^, have emerged as neurochemical signals of interest in modulating taste preference. Studies in humans and animals reported that the neurosteroid allopregnanolone, which acts preferentially at extrasynaptic GABA receptors^15^, induces hyperphagia^16–18^. In humans, allopregnanolone has been associated with binge eating events^16^. Moreover, elevated levels of allopregnanolone have been detected in the blood of obese men and women^16,19^. In rodents, systemic administration of allopregnanolone results in increased intake of highly caloric food independently of palatability^17,20,21^, suggesting an effect of the neurosteroid on taste preference.

Allopregnanolone is a potent modulator of tonic inhibition mediated by GABA extrasynaptic receptors containing the delta subunit (δ-GABA_A_Rs)^14,15,22,23^, widely expressed in cortical and subcortical circuits^22,24^. Activation of δ-GABA_A_Rs engages a slow, long-lasting, inhibitory current^22,25^, suggesting that the effects of allopregnanolone on eating behaviors depend on the modulation of neural circuit excitability. The tonic current is developmentally regulated^26,27^, and, in adults, it represents a larger portion of the total GABA_A_R-mediated current compared to that due to spontaneous synaptic activity^26–29^.

Here, we investigated the role of neurosteroid-sensitive extrasynaptic GABA_A_ receptors in mouse GC. We report that in GC δ-GABA_A_Rs are expressed in subsets of excitatory neurons, and inhibitory parvalbumin- (PV) and somatostatin- (SST) expressing neurons, with the highest expression in SST neurons and lowest excitatory neurons. Bath application of allopregnanolone in acute brain slices elicited slow tonic inhibitory currents and modulated the kinetics of spontaneous inhibitory synaptic currents (sIPSCs), consistent with previous reports in other brain regions. Local infusion of allopregnanolone in GC of male and female mice reduced the sensitivity and preference for sucrose. The effect of allopregnanolone was mediated by GC δ-GABA_A_Rs, as assessed in experiments in which these receptors were knocked down locally within this region. Furthermore, removing δ-GABA_A_Rs selectively from GABAergic neurons in GC was sufficient to abolish the preference for sucrose. Together, our results reveal that δ-GABA_A_Rs in GC are essential for modulating the sensitivity to sweet taste in adult mice and provide the first direct evidence that neurosteroid-sensitive tonic inhibition regulates taste preference, possibly influencing feeding behaviors.

## Results

The distribution and expression patterns of δ-GABA_A_Rs have been examined in cortical and subcortical areas^22,24^, revealing region-specific patterns and suggesting a role for these receptors in different aspects of brain function^30^. There is currently no information regarding the presence and distribution of δ-GABA_A_Rs in GC. We filled this information gap using fluorescent in situ hybridization (FISH, Figure 1A). FISH on brain slices spanning GC revealed mRNA expression, quantified by the presence of one (or more) puncta of δ-GABA_A_R on both GABAergic (co-labeled with probes for Glutamic Acid Decarboxylase, GAD^+^) and putative excitatory neurons (GAD^-^). The labeling was more prominent in GAD^+^ neurons which showed δ-GABA_A_R puncta in 82% of identified GAD^+^ somata, compared to 40% of co-labeled GAD^-^ putative pyramidal neurons (Figures 1B, C). Further analysis on the subsets of neurons showing co-labeling validated the colocalization of δ-GABA_A_R with inhibitory and excitatory neurons, as the overlap coefficients were 0.65 for co-labeled GAD^+^ neurons and 0.60 for co-labeled GAD^-^ neurons; Figure 1D). Analysis of the mean intensities of GAD^+^ and δ-GABA_A_R markers within neuron somata showed a positive correlation (r = 0.62, Figure E) that was lost when one channel was flipped on one of the axes down the z-stack (r = 0.39, Figure F), indicating that the colocalization is not due to random overlap due to tissue processing or imaging artifacts.

**Figure 1.**
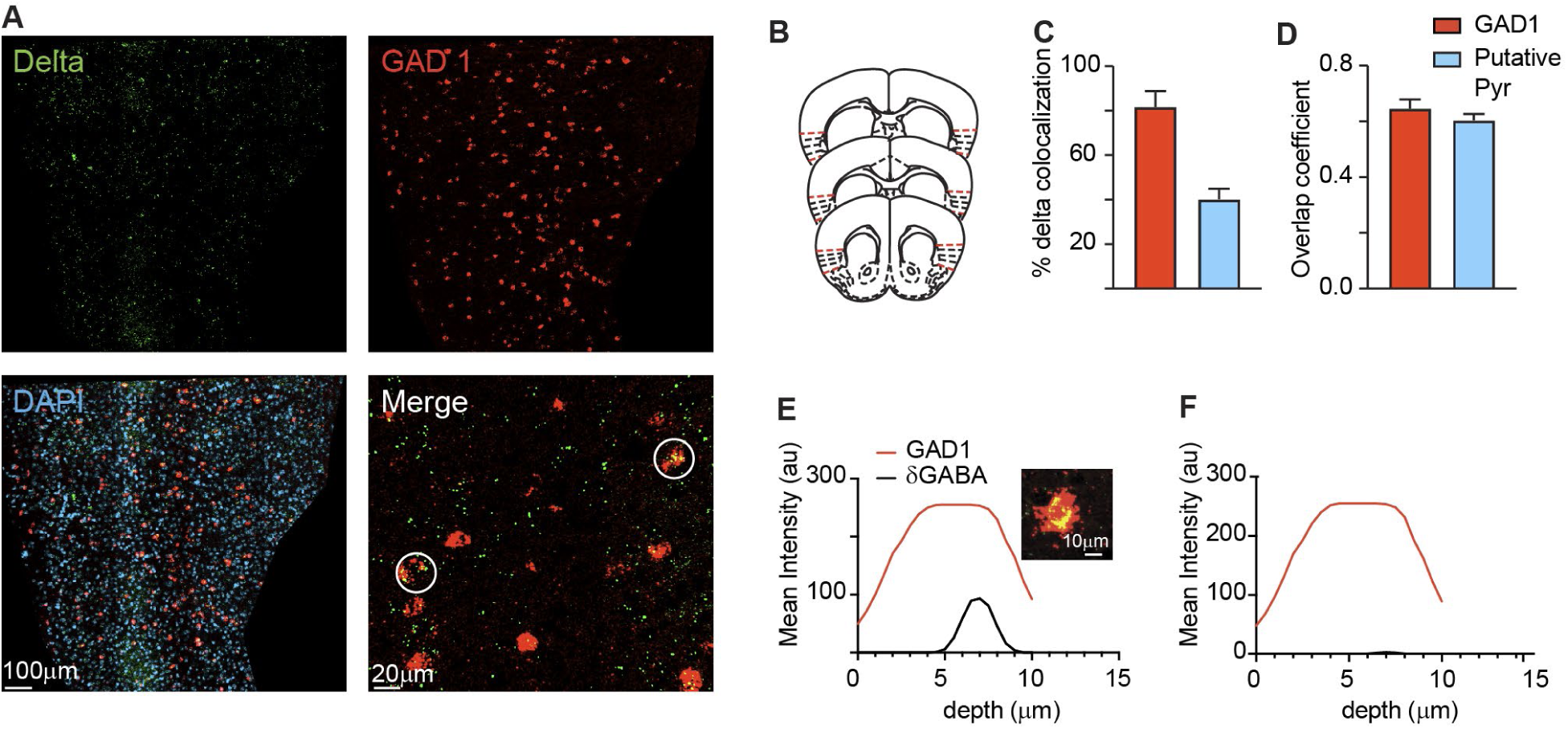
Expression of δ-GABA_A_R subunit in GC using Fluorescence In-Situ Hybridization (FISH). **A.** Localization of δ-GABA_A_R (green) on GAD1 inhibitory neurons (red) and putative excitatory neurons (DAPI-blue) in mouse GC. Co-localized δ-GABA_A_R subunit with GAD1 (merge; yellow). **B.** Coronal GC slices collected for image analysis (from Bregma: +1.53mm to + 0.73mm). **C.** Quantification of δ-GABA_A_R subunit expression. δ-GABA_A_R subunit is expressed in a higher proportion on inhibitory neurons compared to putative excitatory neurons. **D.** Overlap coefficients for δ-GABA_A_R expression on inhibitory and excitatory neurons support validity of co-expression. **E.** Correlation plot of mean intensities of δ-GABA_A_R and example GAD1 cell. **F.** Loss of correlation when δ - GABA_A_R is flipped on the x-axis down the z-stack. Data were obtained from 3 mice (2-3 sections per mouse).

Since other brain regions reported cell type differences in δ-GABA_A_R expression in excitatory neurons as well as somatostatin (SST) and parvalbumin (PV) inhibitory neurons^24^, we used immunohistochemistry to compared expression among putative excitatory neurons and these two inhibitory neuron types (Figure 2A). As observed with FISH, δ-GABA_A_Rs were detected in a larger proportion of inhibitory neurons (∼65%) compared to putative excitatory neurons (32%). When quantified by cell type, the proportion of SST neurons expressing δ-GABA_A_Rs (70%) was significantly larger compared to PV neurons (61%) (Fig. 2B, C). The co-labeling in all three neuron groups was confirmed by analysis of the overlap coefficient (0.70 for δ-GABA_A_R for both SST and PV cells and 0.58 for putative pyramidal cells; Figure 2D). The correlation coefficients for SST (r=0.83) and PV (r=0.72) and the δ-GABA_A_R signal further confirms the colocalization of these markers (Figure 2E, F). To determine whether the colocalization was non-random, we reassessed the correlation coefficients after flipping the images for one channel and found that the correlations were much weaker (Figure 2G, H; SST: r=0.18; PV: r=0.27). Altogether, these results provide evidence of δ-GABA_A_R expression in GC with a preferential association with inhibitory neurons.

**Figure 2.**
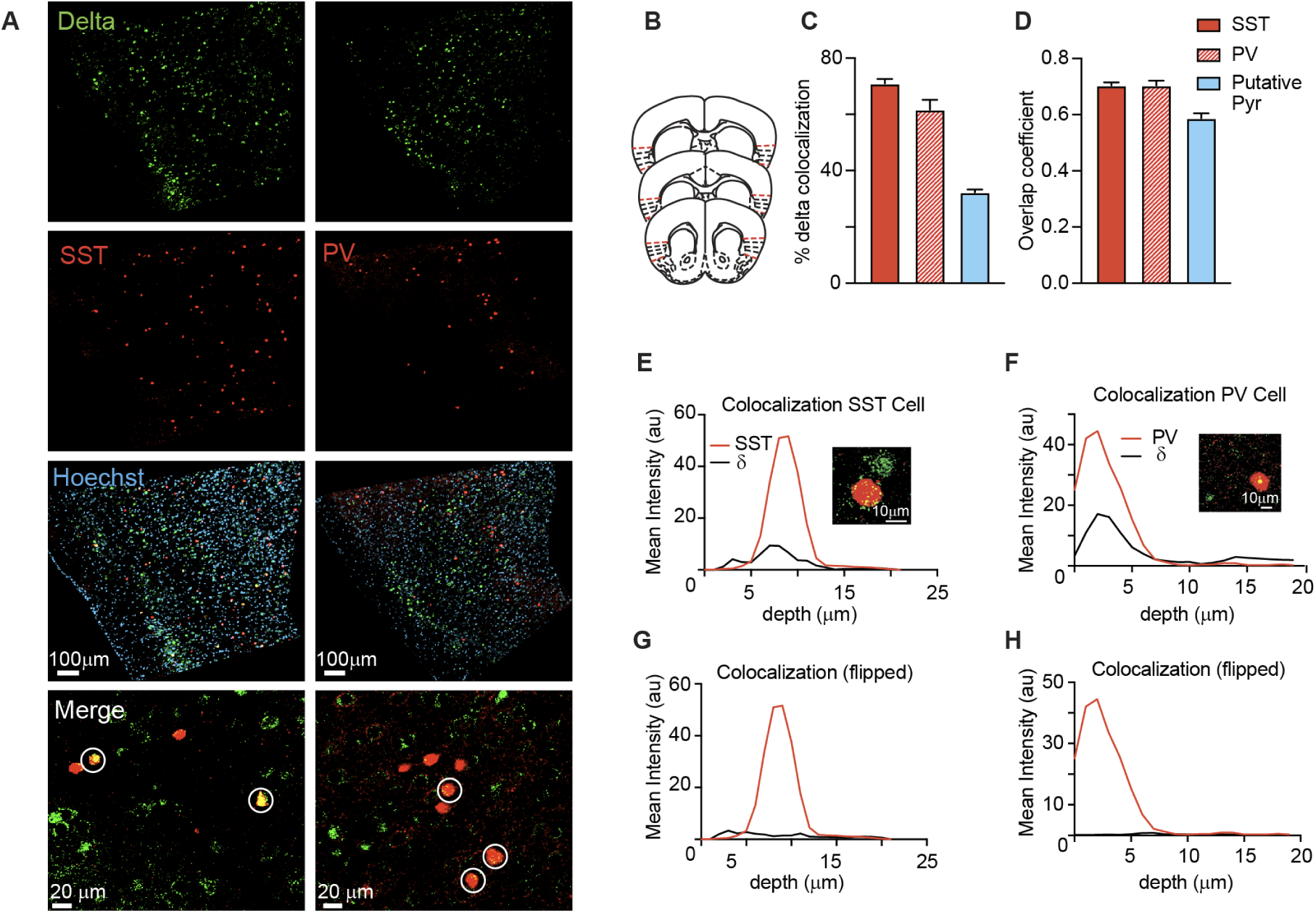
Immunoreactivity of δ-GABA_A_R subunit on SST and PV neurons. **A.** δ-GABA_A_R (green) on SST and PV neurons (red) and putative excitatory neurons (DAPI-blue) in adult mouse GC. δ-GABA_A_R subunit is co-localized with SST and PV cells (merge; yellow). **B.** Coronal GC slices collected for image analysis (from Bregma: +1.53mm to + 0.73mm). **C.** δ-GABA_A_R is expressed in higher proportion on SST neurons, followed by PV and then putative excitatory neurons. **D.** Overlap coefficients for δ-GABA_A_R expression on SST, PV and putative excitatory neurons. **E.** Correlation plot of mean intensities of δ-GABA_A_R puncta and example SST cell. **F.** Correlation plot of mean intensities of δ-GABA_A_R puncta and example PV cell. **G.** Loss of correlation when δ-GABA_A_R is flipped on the x-axis down the z-stack on SST cells. **H.** Loss of correlation when δ-GABA_A_R is flipped on the x-axis down the z-stack on PV cells. Data were obtained from 4 PV-cre and 4 SST-cre mice (3 sections/mouse).

### Allopregnanolone’s modulation of tonic and synaptic inhibition onto GC pyramidal neurons

Tonic inhibitory currents mediated by δ-GABA_A_Rs are strongly modulated by the neurosteroid allopregnanolone (ALLO)^14^. To assess whether δ-GABA_A_Rs quantified anatomically could elicit tonic currents in the presence of ALLO and to measure the tonic current, we obtained patch clamp recordings from visually identified GC neurons in acute slice preparations from adult mice and bath-applied ALLO (500 nM). In Figure 3 we report the effect of ALLO on membrane potential and spontaneous inhibitory synaptic currents (sIPSCs) recorded from excitatory pyramidal neurons in GC. We confirmed the excitatory identity of recorded neurons by post hoc reconstruction of morphology and absence of co-labeling for glutamic acid decarboxylase 67 (GAD67, Figures 3A and 3B). Upon bath application of ALLO, we observed an upward shift in holding current compared to baseline in 36% of recorded pyramidal neurons, as indicated in the representative trace and in the population data in Figure 3C-E. This tonic current was mediated by GABA_A_Rs, as it was abolished by subsequent bath application of picrotoxin (PTX; Figure 3C-E; one-way analysis of variance (ANOVA) *F* _(1.7,14)_ = 12.07, *P* = 0.001, with post hoc Tukey corrected tests: ACSF versus ALLO, *P* = 0.023; ACSF versus PTX, *P* = 0.166; and ALLO versus PTX, *P* = 0.005). As suggested by FISH and immunohistochemistry data indicating that only a percentage of putative pyramidal neurons express δ-GABA_A_Rs, a substantial proportion of recorded excitatory neurons (64%) showed no change holding current upon bath application of ALLO (Figure 3F-H; Friedman Test χ^2^(3) = 15, *P* = 0.0002, with post hoc Dunn’s tests: ACSF versus ALLO, *P* = 0.231; ACSF versus PTX, *P* = 0.0001; and ALLO versus PTX, *P* = 0.0647). Previous studies reported ALLO modulation of synaptic currents in addition to tonic currents^31–33^. To assess whether GC synaptic inhibition was also affected, we quantified amplitude, kinetics and frequency of sIPSCs in neurons with and without a tonic current. In neurons showing an increased tonic current following ALLO application (Figure 3I), we observed no changes in sIPSC amplitude (Figure 3J; two-tailed paired t-test, *P* = 0.094) but increased rise time (Figure 3K; two-tailed paired t-test, *P* = 0.008) and decay time (Fig. 3L; two-tailed paired t-test, *P* = 0.005), pointing to an increased duration of synaptic inhibitory currents.. The frequency of sIPSCs was significantly decreased (Figure 3M; two-tailed paired t-test, *P* = 0.003), suggesting presynaptic modulation of inhibitory synapses in addition to changes in postsynaptic currents. In excitatory neurons without a shift in holding current (Figure 3N), following ALLO application we observe a decrease in sIPSC amplitude (Figure 3O; two-tailed paired Wilcoxon test, *P* = 0.039) and no changes in sIPSC kinetics, (rise time: Figure 3P; two-tailed paired Wilcoxon test, *P* = 0.495; decay time: Figure 3Q; two-tailed paired t-test, *P* = 0.053), nor frequency (Figure 3R; Wilcoxon Signed-Rank Test, *P* = 0.562). These results suggest differential modulation by ALLO in pyramidal neurons expressing δ-GABA_A_Rs-mediated tonic current and those that do not.

**Figure 3.**
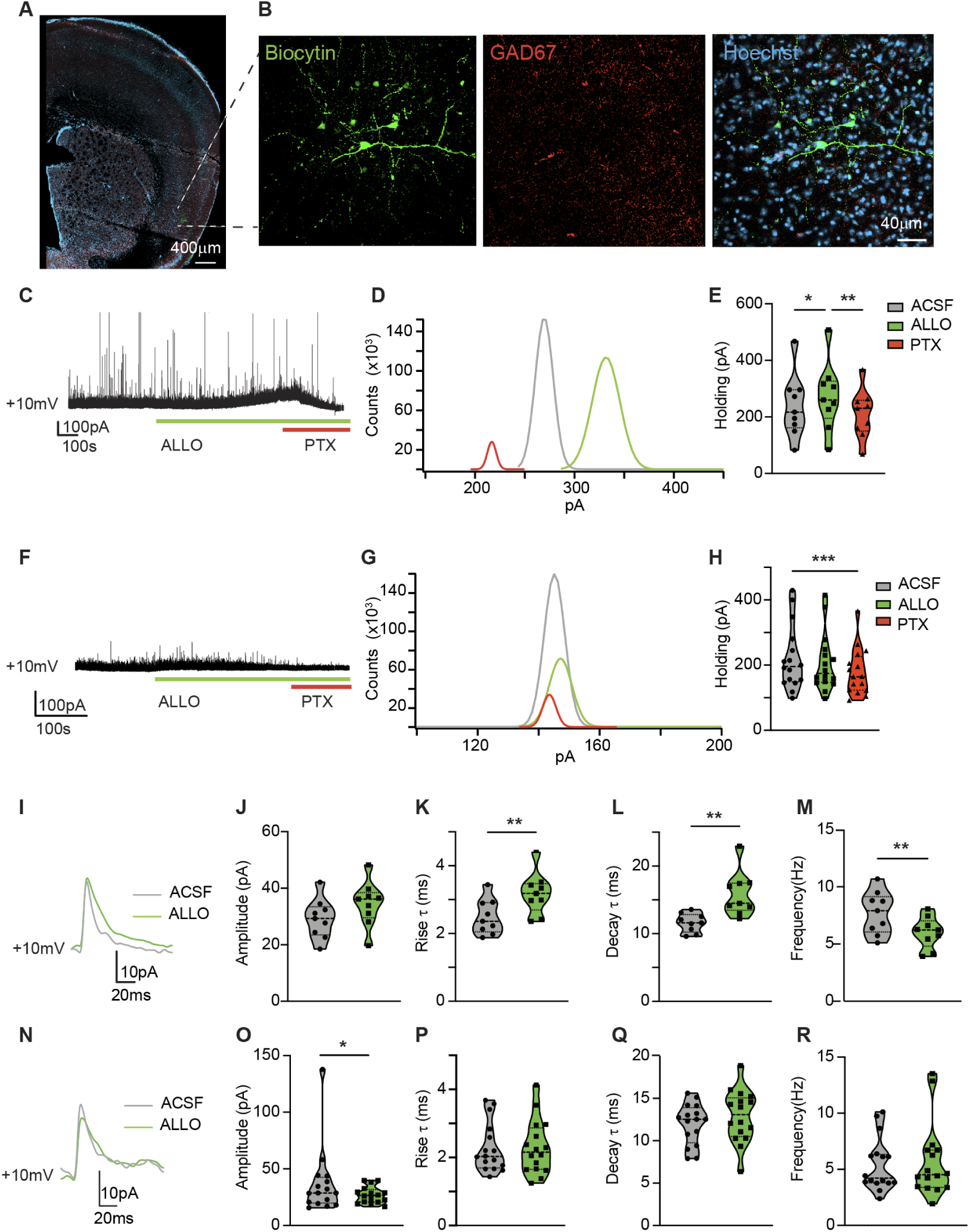
Allopregnanolone modulates inhibitory synaptic transmission on excitatory neurons in GC. **A.** Representative image of a recorded pyramidal neuron in a brain slice containing GC. **B.** Example of a recorded GC excitatory neuron showing pyramidal morphology and no co-labeling of GAD67 (Green, biocytin-filled neuron; Red, GAD67 immunostaining; Cyan, Hoechst nuclei counterstain). **C.** Representative pyramidal cell trace showing increased tonic current in a pyramidal neuron. This effect was observed in a proportion of recorded pyramidal neurons (n = 9) **D.** All-points histogram depicts positive shift in tonic current. **E.** Violin plot summarizing change in holding current following ALLO application. **F.** Sample pyramidal cell trace showing no effect of ALLO on tonic current in a percentage of pyramidal neurons. This effect was observed in a proportion of pyramidal neurons (n = 16). **G.** All-points histograms confirm no change in tonic current. **H.** Violin plot summarizing change in holding current following ALLO application and following picrotoxin. **I.** Representative overlay of average sIPSC from a pyramidal cell showing ALLO-dependent tonic current before (gray) and after ALLO (green). Note the slower kinetics of sIPSCs after ALLO application. **J-M.** Violin plots show no change in sIPSC event amplitude (J), significant increases in rise time (K) and increase in decay time (L) and increased frequency (M). **N-R.** Representative overlay of average sIPSC events from a pyramidal cell with non-altered tonic current pre (gray) and post ALLO (green) (N). Violin plots show increase in event amplitude (O), no change in rise time (P) and decay time (Q) and no change in frequency (R). Data were obtained from N = 10 mice; * = P ≤ 0.05, ** = P ≤ 0.01, *** = P≤ 0.01.

### Effect of allopregnanolone on tonic and synaptic currents onto GC inhibitory neurons

Our histology data show that a large proportion of inhibitory neurons in GC express, δ-GABA_A_Rs. Here, we asked whether bath application of ALLO can evoke an increase in holding current, which is typical of the activation of tonic inhibition. We recorded SST neurons as identified by tdTomato-expressing cells in GC of *SST-Cre;Ai14* mice and confirmed the cell type with post hoc immunohistochemistry to identify co-labeling of biocytin and tdTomato (Figure 4A-C). In 46% of recorded SST neurons bath application of ALLO induced an upward shift in holding current compared to baseline, as indicated in representative traces and population data. The effect of ALLO was abolished by PTX (Figure 4D-F; Friedman Test χ^2^(3) = 9.3, *P* =0.006, with post hoc Dunn’s tests: ACSF versus ALLO, *P* = 0.021; ACSF versus PTX, *P* = 0.564; and ALLO versus PTX, *P* = 0.004). As in pyramidal neurons, a subset of SST neurons (54%) showed no shift in holding current during ALLO application (Figure 4G-I; (ANOVA) *F* _(1.1,7.9)_ = 5.9, *P* = 0.039, with post hoc Tukey’s tests: ACSF condition versus ALLO condition, *P* = 0.730; ACSF versus PTX, P = 0.089; and ALLO versus PTX, *P* = 0.105). When we assessed possible modulation of sIPSCs in SST neurons showing an ALLO-induced increase in tonic inhibitory current, we observed no changes in sIPSC amplitude, rise and decay kinetics (Figure 4J), and frequency (amplitude: Figure 4K; two-tailed paired Wilcoxon test, *P* > 0.999; rise time: Figure 4L, two-tailed paired t-test, *P* = 0.931; decay time: Figure 4M, two-tailed paired t-test, *P* = 0.446; frequency: Figure 4N, two-tailed paired t-test, *P* = 0.844). SST neurons with no shift in holding current following ALLO showed no changes in sIPSC amplitude and rise kinetics (Figure 4O; amplitude: Figure 4P, two-tailed paired Wilcoxon test, *P* = 0.688; rise time: Figure 4Q, two-tailed paired t-test, *P* = 0.334). However, we observed an increase in decay time constant (Figure 4R; two-tailed paired Wilcoxon test, *P* = 0.016), suggesting possible modulation of GABA channel opening time, and no change in frequency (Figure 4S, two-tailed paired Wilcoxon test, *P* = 0.469).

**Figure 4.**
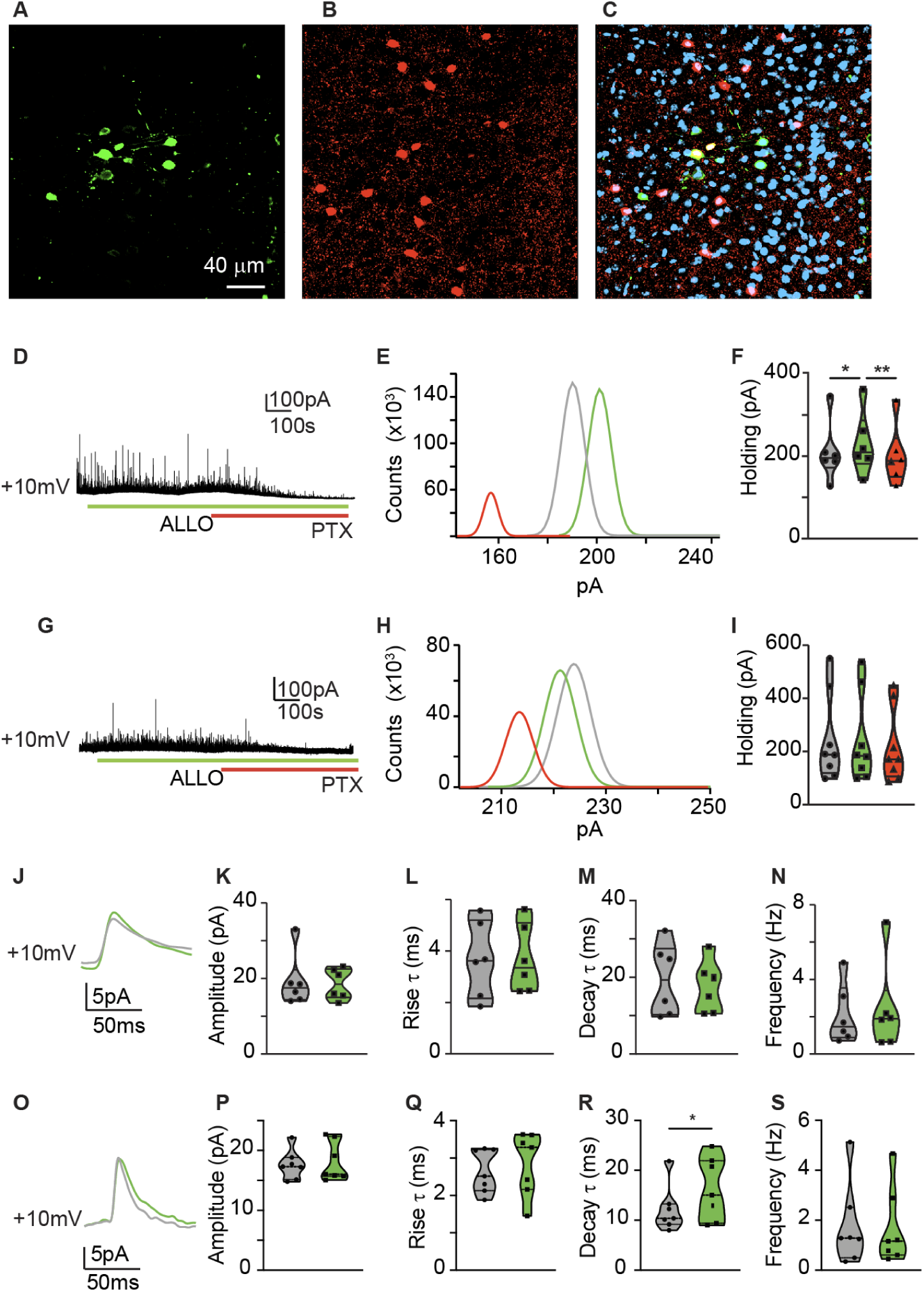
Allopregnanolone modulates inhibitory synaptic transmission on inhibitory SST neurons in GC. **A.** Recorded biocytin-filled (Green) neuron. **B.** tdTomato expression in SST positive neurons (red). **C.** Image of recorded cell co-expressed with SST neuron in GC (Cyan, Hoechst nuclei counterstain). **D.** Representative trace showing increased tonic current in response to bath application of ALLO. This effect was observed in a proportion of recorded SST neurons (n = 6). **E.** All-points histogram depicts increase in tonic current. **F.** Violin plot summarizing change in holding current. **G.** Example trace showing lack of tonic current in a subset of SST neurons following ALLO application in a subset of recorded neurons (n = 7). **H.** All-points histogram of tonic current before and after ALLO and its elimination by PTX. **I.** Violin plot summarizing change in holding current. **J.** Representative average of sIPSC events from a tonic current expressing SST neuron before (gray) and after ALLO (green). **K-N.** Violin plots show no changes in IPSC amplitude (K), rise time (L), decay time (M) and frequency (N) of IPSCs. **O.** Overlay of average sIPSC events from an SST neuron without the tonic current before (gray) and after ALLO (green). **P-S.** Violin plots show no effect of ALLO on sIPSC’s amplitude (P), rise time (Q), decay time (R) and frequency (S). Data were obtained from N = 5 mice; * = P ≤ 0.05, ** = P ≤ 0.01.

Next, we assessed the effect of ALLO on PV neurons. To identify PV neurons we obtained slices from *PV-Cre;Ai14* mice and confirmed recordings with post hoc immunohistochemistry (Figure 5A-C). Application of ALLO induced an upward shift in holding current in 46% of recorded PV neurons (Figure 5D-F; Friedman Test χ^2^(3) = 10, *P* =0.0002, with post hoc Dunn’s tests: ACSF versus ALLO, *P* = 0.043; ACSF versus PTX, *P* = 0.248; and ALLO versus PTX, *P* = 0.002). A subset of PV (54%) showed no change in tonic current following ALLO application (Figure 5G-I; (ANOVA) *F* _(1.5,11)_ = 20, *P* = 0.0004, with post hoc Tukey’s tests: ACSF condition versus ALLO condition, *P* = 0.445; ACSF versus PTX, P = 0.002; and ALLO versus PTX, *P* = 0.01). Further analysis of sIPSCs in PV neurons showing an ALLO-induced shift in tonic inhibitory current showed no changes in sIPSC amplitude, rise and decay kinetics (Figure 5J) and an increase in frequency (amplitude: Figure 5K; two-tailed paired t-test, *P* = 0.461; rise time: Figure 5L, two-tailed paired t-test, *P* = 0.285; decay time: Figure 5M, two-tailed paired t-test, *P* = 0.548; frequency: Figure 5N, two-tailed paired t-test, *P* = 0.046). PV neurons with unaltered holding current following ALLO showed no changes in sIPSC amplitude, rise and decay kinetics (Figure 5O) and frequency (amplitude: Figure 5P, two-tailed paired t-test, *P* = 0.177; rise time: Figure 5Q, two-tailed paired Wilcoxon test, *P* = 0.469; decay: Figure 5R, two-tailed paired t-test, *P* = 0.246; frequency: Fig. 5S, two-tailed paired Wilcoxon test, *P* = 0.874).

**Figure 5.**
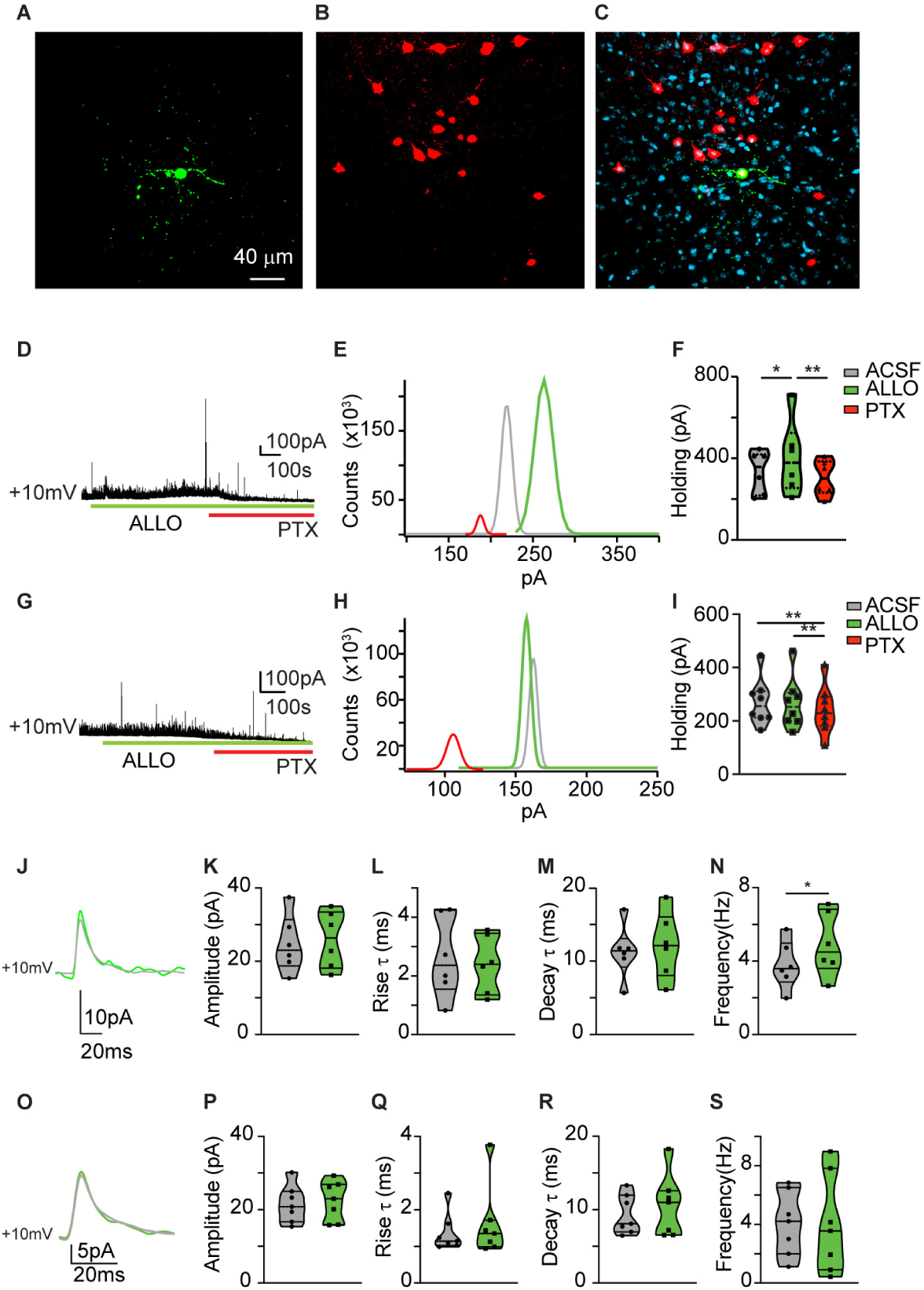
Allopregnanolone modulates inhibitory synaptic transmission on inhibitory PV neurons in GC. **A.** Recorded biocytin-filled (Green) neuron. **B.** tdTomato expression in PV positive neurons (red). **C.** Merged image of recorded cell with co-expression with PV neuron in GC (Cyan, Hoechst nuclei counterstain). **D.** Representative trace showing ALLO-dependent tonic current in a proportion of PV neurons (n = 6). **E.** All-points histogram reporting positive shift in tonic current and their elimination by picrotoxin. **F.** Violin plot summarizing changes in holding current. **G.** Example trace showing unchanged tonic current with ALLO application. This effect was observed in a subset of PV neurons (n = 7). **H.** All-points histogram showing unaltered tonic current. **I.** Violin plot summarizing change in holding current. **J.** Representative average of sIPSC events from a PV neuron expressing a tonic current recorded before (gray) and after ALLO (green). **K-M.** Violin plots show no change in phasic event amplitude (K), rise time (L) and decay time (M). **N.** ALLO increased the frequency of sIPSCs in PV neurons expressing the tonic current. **O.** Overlay of average sIPSC events from a PV neuron without the tonic current recorded before (grey) and after ALLO (green). **P-S.** Violin plots show no effect of ALLO on sIPSC’s amplitude (P), rise time (Q), decay time (R) and frequency (S). Data were obtained from N = 5 mice; * = P ≤ 0.05, ** = P ≤ 0.01.

These results indicate that ALLO modulates inhibitory drive onto distinct neuron types in GC. ALLO increases tonic inhibition, possibly by directly activating neurosteroid-sensitive receptors like δ-GABA_A_Rs. ALLO also shows modulation of sIPSCs, which is consistent with reports of increased tonic currents and altered sIPSCs kinetics in other circuits^28,32,34,35^.

### Localized infusion of allopregnanolone modulates taste preference

The GC is involved in taste perception, palatability and preference^3,5,7,8^. As ALLO activates tonic inhibitory currents onto GC excitatory and inhibitory neurons, it may modulate taste processing. We asked whether local infusion of ALLO in GC may affect the preference for sweet taste in adult mice. To assess taste preference, we trained mice to perform a brief access test (BAT) (Figure 6A)^8,36^. Adult WT mice bilaterally implanted with infusion cannulae were exposed to water (0 mM sucrose) and 4 concentrations of sucrose (in mM: 10, 50, 200, 600) for multiple 10 s long self-initiated trials in a Davis Rig gustometer (Figure 6A)^36^. Sucrose preference was assessed by normalizing the number of licks for each sucrose concentration to that of water (tastant-water (T/W) lick ratio)^8,36^ following infusion of saline (control) or 500 nM ALLO locally in GC. Only mice with histologically verified position of cannulae in GC were included in the analysis (Figure 6B). Infusion of ALLO significantly reduced the total number of licks per experimental session (Figure 6C; two-tailed paired t-test, *P* = 0.015). We also observed a reduction in the number of self-initiated trials (Figure 6D; two-tailed paired t-test, *P* = 0.003). Further examination of BAT results indicated a reduction in sucrose preference following the infusion of ALLO in GC (Figure 6E). A two-way ANOVA was used to examine concentration and drug effects on preference. There was a significant main effect of sucrose concentration on sucrose preference, F (2.1, 10) = 23, P=0.0001. There was also a significant main effect of ALLO, F (1.0, 5.0) = 41, P=0.001. The interaction between sucrose concentration and drug was not significant, F (1.3, 6.6) = 3.6, P=0.099. Post hoc comparisons adjusted for multiple testing using Šidák’s multiple comparisons test indicate that the mean lick number following ALLO infusion was significantly lower at 50 mM (with a mean difference of 0.32, SE = 0.073, and an adjusted p-value of 0.037) and 200 mM concentrations (mean difference of 0.42, SE=0.094, and adjusted p-value of 0.033) compared to control. The mean scores for other concentrations did not differ significantly (Water, P > 0.999; 10 mM, P = 0.989; 600 mM, P = 0.235). The normalized licks were plotted against sucrose concentration to generate a preference curve, and a function was fit to each animal’s sucrose curve generated following saline and the curve generated following ALLO infusion with a sigmoidal 3-parameter logistic function [Y=b+((a−b)/ 1+10^(logEC50−X)^)] to determine whether a single curve was sufficient to fit both conditions^8^. We found that two separate curves were needed, indicating a difference in the sucrose/water lick ratio at concentrations equal or higher than 50 mM. Furthermore, the EC50 significantly increased following ALLO, pointing to a rightward shift in the sucrose curve (saline = 96.67, ALLO = 101.5, F_(3,54)_ = 4.661, P = 0.0057), an effect that indicates decreased sensitivity to sucrose^37^. Thus, local infusion of ALLO in GC reduced the preference for sucrose by shifting its sensitivity to higher concentrations^37^ and decreasing the T/W lick ratio^8^.

**Figure 6.**
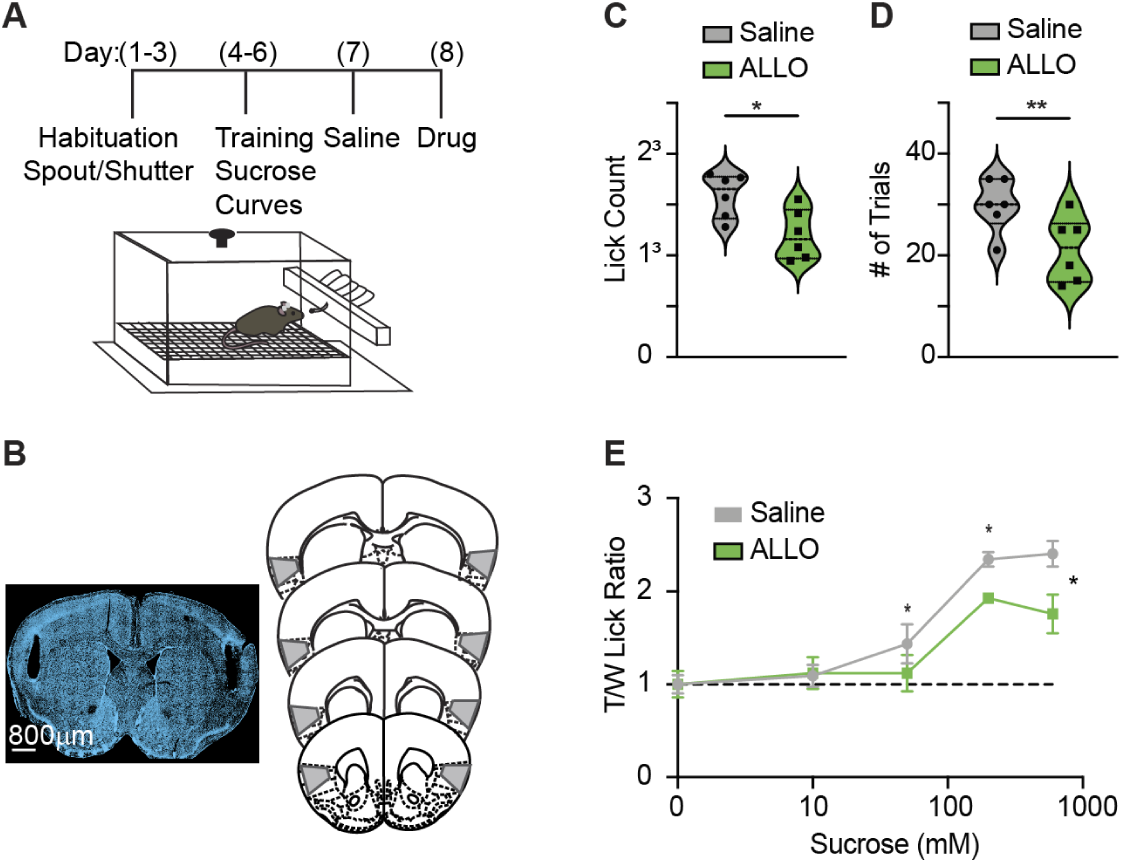
Allopregnanolone alters taste sensitivity and sucrose preference in WT mice. **A.** Top: Sucrose preference test schedule. Bottom: Representation of Davis rig gustometer. **B.** Representative cannulae placements in GC and registration to mouse brain atlas. **C.** Lick counts following saline (grey) and after ALLO (green) infusion. **D.** Number of trials following saline (grey) and ALLO (green) infusions. **F.** Sucrose preference curves for WT mice following saline (grey) and ALLO (green) infusion. Data were obtained from N = 6 mice; * = P ≤ 0.05, ** = P ≤ 0.01.

### δ-GABA_A_Rs mediate the modulation of sucrose preference by allopregnanolone

To determine whether the effect of ALLO on sucrose preference engaged δ-GABA_A_Rs, we took advantage of genetic manipulations. We used floxed Gabrd mice^29^, which allow for the deletion of exons 2 through 5 and a portion of exon 6 of the delta gene with cre-dependent constructs, making δ-GABA_A_Rs nonfunctional (Figure 7A). The insular cortex is an epileptogenic region^38^, complete elimination of tonic inhibition in homozygous floxed Gabrad mice may lead to hyperexcitability of the circuit^39^ and affect behavior. To avoid this potential for instability, we performed the following experiments in heterozygotes floxed Gabrd mice. In a group of animals, we used a viral construct that allowed for manipulating δ-GABA_A_Rs activity in GC independently of a neuron’s excitatory or inhibitory identity (AAV9-Cre-GFP) to assess the overall contribution of tonic inhibition to sucrose taste preference. Following bilateral viral injection localized in GC, experimental mice were implanted with infusion cannulae in GC and tested on the BAT to assess sucrose preference (Figure 7A). Injection sites and cannulae placements were verified postmortem before further analysis (Figure 7B). Compared to WT animals, Cre-GFP mice showed a significantly reduced preference for sucrose especially at higher concentrations (Figure 7C; 2-way ANOVA Mixed effects: Concentration effect, P <0.0001; Mutant Effect, P <0.0001, Concentration x Mutant effect: P = 0.043; Šidák’s multiple comparisons test: Water, P > 0.999; 10mM, P = 0.625; 50mM, P = 0.174; 200mM P = 0.001; 600mM, P = 0.003). The shift was further quantified as a significant change in the EC50 (WT = 96.67, Cre-GFP = 189.7; F_(3,54)_ = 11.6, P < 0.001), suggesting that loss of δ-GABA_A_Rs in excitatory and inhibitory decreased sensitivity to sucrose. This effect was unexpected, as it shows how removal of δ-GABA_A_Rs has a similar effect to activation of tonic inhibition by ALLO. A possible interpretation of this findings is that the the effects of ALLO infusion does not depend on functional δ-GABA_A_Rs. If this is the case, infusion of ALLO in mice with δ-GABA_A_Rs knock down should modulate sucrose preference further. To test this, in a subset of cre-GFP mice we infused saline or ALLO in GC before testing them on BAT. In mice with knock down of δ-GABA_A_Rs in excitatory and inhibitory neurons the effect of ALLO was completely occluded as there was no significant change in the sucrose curve (Figure 7D; 2-way ANOVA Repeated Measures: Concentration effect, P = 0.004; Drug Effect, P = 0.064, Concentration x Drug effect: P = 0.449; Šidák’s multiple comparisons test: Water, P > 0.999; 10mM, P > 0.999; 50mM, P = 0.814; 200mM P = 0.123; 600mM, P = 0.691). This finding confirms that ALLO’s effect on the sucrose curve was mediated by δ-GABA_A_Rs in GC and suggests that modulation of tonic inhibition is involved in maintaining the preference for sucrose.

**Figure 7.**
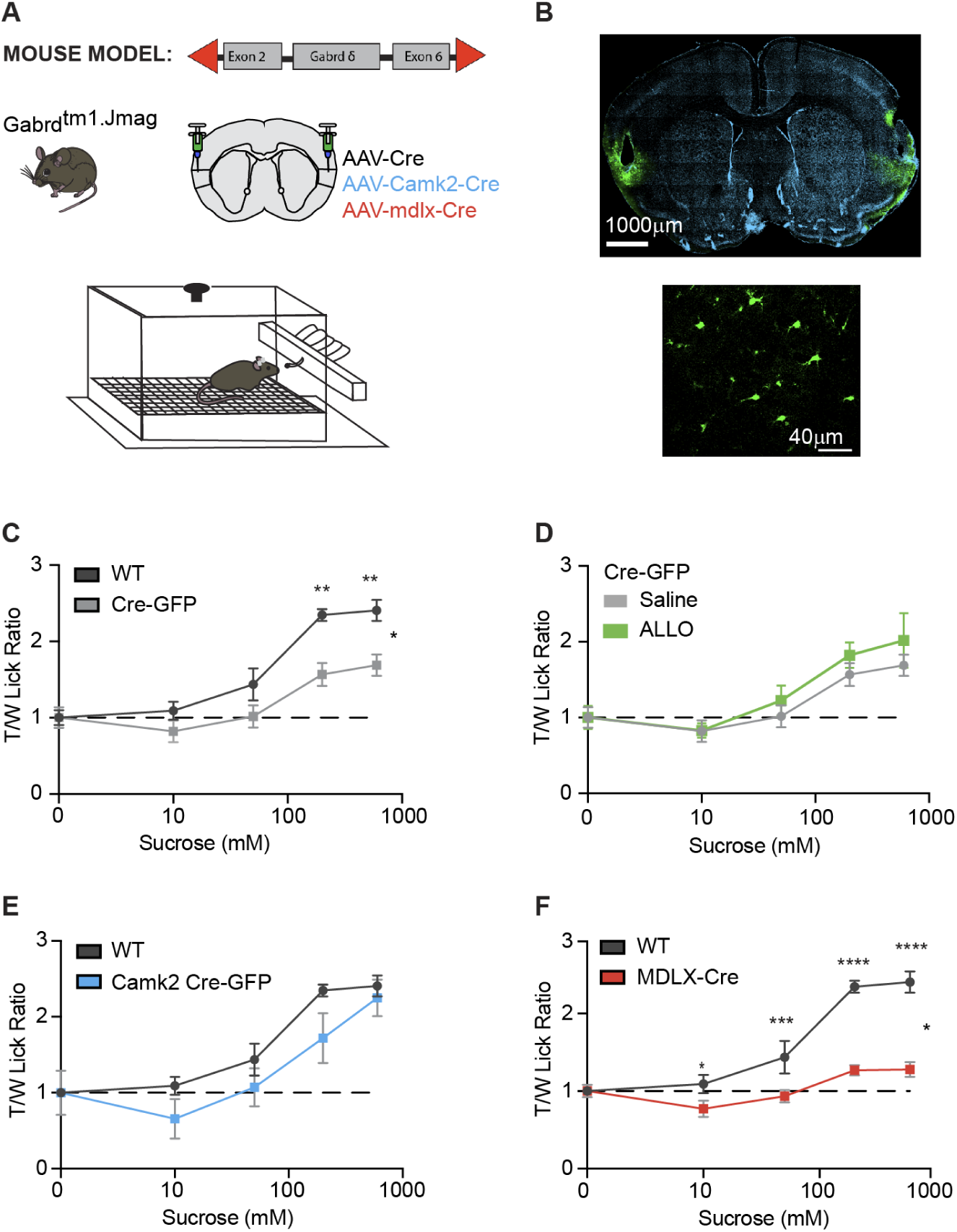
Conditional knockout of δ-GABA_A_Rs in GC alters taste sensitivity and sucrose preference in mice. **A.** Top. Cartoon of Gabrd mouse model with floxed delta gene and diagram of viral injection targeting, and viral constructs. Bottom. Representation of Davis rig gustometer. **B.** Top. Representative cannulae placement in GC and histological verification of injection sites. Bottom. Zoomed GC image showing GFP-expressing neurons indicating knock down of δ-GABA_A_Rs. **C.** Sucrose curve for WT (controls; N = 6 mice; black) and floxed Gabrd mice with Cre-GFP injections (N = XX mice; grey). **D.** Sucrose curve for Gabrd;Cre-GFP mice following saline (grey) and ALLO (green) infusion. **E.** Sucrose curve for WT controls (black, same curve as in C) and floxed Gabrd mice with CamKII Cre-GFP injections (N = 6 mice; blue). **F.** Sucrose curve for WT controls (black, same curve as in C) and floxed Gabrd mice with mDLX Cre-RFP injection (N = 6 mice, red). * = P ≤ 0.05, ** = P ≤ 0.01, *** = P≤ 0.01.

In view of the differential expression of δ-GABA_A_Rs in excitatory and inhibitory neurons, we then asked whether the behavioral effect of ALLO may depend on the neurochemical identity of the neurons expressing δ-GABA_A_Rs. To test this possibility, we knocked down δ-GABA_A_Rs function selectively in excitatory neurons (AAV9-CAMKII Cre-GFP; Figure 7E) or inhibitory neurons (AAV9-mDLx Cre-RFP; Figure 7F) in separate groups of heterozygous floxed Gabrd mice. In mice in which δ-GABA_A_Rs was knocked down selectively in pyramidal neurons (CAMKII Cre-GFP), although the curve appeared shifted and there was no preference for sucrose over water at 10 mM and 50 mM, the difference in the curve was not statistically significant and the curves were fit by the same logistic function (Figure 7E; 2-way ANOVA Mixed effects: Concentration effect, P <0.0001; Mutant Effect, P <0.122, Concentration x Mutant effect: P = 0.454; Šidák’s multiple comparisons test: Water, P > 0.999; 10 mM, P = 0.596; 50 mM, P = 0.824; 200 mM, P = 0.558; 600 mM, P = 0.995). There were also no significant differences in the slope of the curves nor in the EC50 (WT = 96.67, CAMKII Cre-GFP = 138.5, F_(3,49)_ = 2.485, P < 0.072). These results suggest a lack of a significant modulation of the sucrose curve in mice in which δ-GABA_A_Rs were selectively knocked down in GC pyramidal neurons.

Differently, mice in which δ-GABA_A_Rs function was knocked down selectively in GABAergic neurons (mDLx Cre-RFP) showed a flattened sucrose curve, indicating that preference for sucrose over water was practically abolished (Figure 7F; 2-way ANOVA Mixed effects: Concentration effect, P <0.0001; Mutant Effect, P <0.0001, Concentration x Mutant effect: P <0.0001; Šidák’s multiple comparisons test: Water, P > 0.999; 10mM, P = 0.047; 50mM, P = 0.001; 200mM P < 0.0001; 600mM, P <0.0001). In addition, there was a significant change in fitted slope and EC50 (WT = 96.67, mDLx Cre-GFP = 180.1, F_(3,54)_ = 32.55, P <0.0001). These data demonstrate that the effect of ALLO on sucrose preference depends on δ-GABA_A_Rs in GC. The effect of activation of δ-GABA_A_Rs by ALLO and the knock down of δ-GABA_A_Rs in GC have comparable effects on sucrose preference suggesting that sucrose preference depends on a balance in the modulation of tonic currents mediated by δ-GABA_A_Rs. Finally, we show that δ-GABA_A_Rs on inhibitory neurons are essential for preserving any sucrose sensitivity and preference in adult mice.

## Discussion

We report that neurosteroid-mediated tonic GABAergic currents in mouse GC modulate sensitivity and preference for sweet taste in adult mice. The action of the neurosteroid allopregnanolone relied on the presence of δ-GABA_A_Rs on subsets of GC inhibitory and excitatory neurons. The cell types expressing δ-GABA_A_Rs are consistent with previous reports from other brain regions, including thalamus, somatosensory cortex and hippocampus^22,35,40–43^. However, the GC’s expression pattern differs from other areas, where δ-GABA_A_Rs were most highly expressed in PV neurons^24,32,34^, with much lower presence in SST neurons^26,28^. In GC, δ-GABA_A_Rs expression was highest in SST neurons, followed by PV neurons and, to a lesser extent, by pyramidal neurons. These results suggest that neurosteroid modulation of δ-GABA_A_Rs in GC engages more diverse cell types and may affect circuit excitability more broadly and distinctly than other brain regions. Previous studies reported selective modulation of tonic inhibition by neurosteroids^25,44^, while others reported a combination of effects on tonic and synaptic currents^15,31,45,46^. Our data are consistent with findings that neurosteroid modulation of inhibitory drive in GC is not restricted to extrasynaptic receptor activation. While the most prominent effect of allopregnanolone was the activation of a tonic inhibitory current, we also observed subtle modulation of the kinetics of spontaneous IPSCs. The increase in the rise and decay time of sIPSCs points to slower activation and inactivation of inhibitory postsynaptic receptors, consistent with the reported allosteric effect of allopregnanolone, which is thought to increase the duration of GABA synaptic transmission^15,31,46^. While recent work suggests that allopregnanolone may affect GABA_A_Rs containing the γ2 subunit^31,33,46^, typically found at inhibitory synapses^47,48^, our results showing nearly complete occlusion of the effects of allopregnanolone in mice with δ-GABA_A_Rs knockdown hint that the behavioral effects we report are driven predominantly by δ-GABA_A_Rs. WT mice infused with allopregnanolone showed reduced sucrose preference. This result may appear to be in contrast with studies reporting hyperphagic effects of systemic allopregnanolone administration in rodents and humans^17–19,21,49^. However, research on obesity driven by overconsumption reports that alterations in taste perception worsen palatability-induced hyperphagia^50–53^. Studies in obese rats showed shifted sucrose preference towards higher concentrations, suggesting decreased sensitivity for sweet taste^54–56^. In addition, humans with reduced sensitivity to sweet taste were shown to eat more sweets and other highly palatable food^51,57^. These proclivities are attributed to a higher tastant detection threshold, consistent with impaired taste function^57^. In our experiments, allopregnanolone results in a rightward shift of the sucrose preference curve and an increase in the sucrose preference threshold, determined by the increase in EC50 values. We also report a flattening of the sucrose curve, indicating reduced preference. The reduced sucrose sensitivity and the decrease in sucrose preference we report align with numerous studies that have linked decreased taste sensitivity and altered taste perception with changes in food consumption.

Removing δ-GABA_A_Rs from GC neurons decreased sensitivity to sucrose at low concentrations and reduced sucrose preference. Selective removal of δ-GABA_A_Rs from pyramidal neurons reduced sensitivity to sucrose but still showed a preference for sweet over water. Differently, knockdown of δ-GABA_A_Rs from GABAergic neurons not only decreased sensitivity at low concentrations but completely flattened the sucrose curve, suggesting a fundamental role for these receptors in the ability to differentiate a sweet taste from water. The knockdown of δ-GABA_A_Rs in GC neurons occluded any additional effect of allopregnanolone, supporting the interpretation that the effect of allopregnanolone are indeed mediated by modulation of these receptors.

In many cortical regions, including GC, inhibitory neurons shape how excitatory neurons integrate information by sharpening the tuning of pyramidal neurons^58^. Most studies to date focused on the role of specific inhibitory neuron groups in modulating gain or breadth of tuning of pyramidal neurons^58–62^; however, little attention has been given to the contribution of tonic inhibition^29,35^. Our findings underscore the requirement for a fine-tuned modulation of inhibitory drive for taste perception, as we report that both potentiation of tonic currents by allopregnanolone or elimination of tonic inhibition by knockdown of δ-GABA_A_Rs alter the sensitivity and preference of sweet taste. Furthermore, the magnitude of the effect of tonic inhibition on taste depends on the engagement of either excitatory or inhibitory neurons. While ours is the first study looking at the contribution of neurosteroid-sensitive tonic inhibitory currents on taste sensitivity and preference, comparable fine-tuning of modulation has been observed for other neuromodulators such as dopamine and norepinephrine for which inverted U-shaped dose-response curves have been reported^63,64^. Within a sensitive working range, tonic inhibition may help preserve sensitivity to sucrose concentrations, thereby maintaining the innate preference for sweetness that is typically observed in humans and animals. High levels of neurosteroids broadly increase tonic inhibition on distinct neuron types, especially GABA neurons, which may lead to disinhibition of the GC circuit and impaired ability to differentiate sucrose concentrations from water, resulting in diminished sensitivity. A similar effect, though driven by a shift in excitability in the opposite direction, may be induced by deletion of the δ-GABA_A_Rs. As δ-GABA_A_Rs are more highly expressed on SST and PV neurons, loss of tonic inhibition onto these neuron groups may result in their hyperactivation, leading to excessive suppression of pyramidal neurons’ excitability and decreased responsiveness to sucrose concentrations, leading to reduced preference.

Allopregnanolone has been linked to increased food consumption, particularly of energy-dense and palatable foods^17–19,21^. Its dysregulation is implicated in eating disorders, especially binge eating disorders, underscoring its crucial role in the neurobiology of feeding^19,49^. Our results, which demonstrate that acute allopregnanolone signaling alters the sensitivity and preference for sweet taste through the activation of tonic inhibitory currents in GC, broaden our understanding of the neurochemical signals for taste perception and their influence on food choice and feeding.

## Acknowledgments

This work was supported by National Institutes of Health grant R01DC019827 to AM

National Institutes of Health grant R01DC013770 to AM and AF

National Institutes of Health grant R01DC015234 to AF and AM

National Institutes of Health grant UF1NS115779 to AF and AM

National Institutes of Health grant F31 DC019518 to PEY

We wish to thank Dr. Jamie Maguire for kindly providing colony founder breeders for the floxed Gabrd mice used in this study and Aylar Berenji-Kalkhoran for assisting with colony maintenance.

## Author Contributions

Designed experiments: PEY, AF, AM

Performed experiments: PEY

Analyzed data: PEY

Supervised experiments and analysis: AF, AM

Writing: PEY, AF, AM

## Materials & Methods

### Animals

All experiments and surgical procedures were approved by the Stony Brook University IACUC and followed the National Institute of Health guidelines. Mice of both sexes were housed in a vivarium on a 12-hour light-dark cycle in standard cages with ad libitum access to food and water (unless otherwise noted). Mice were single-housed following surgeries and during behavioral training. Wild-type (WT) C57BL/6J mice (catalog #000664), *PV-Cre* (catalog #017320), SST-Cre (catalog #013044), and *Ai14* (RRID: IMSR_JAX:007908) were obtained from The Jackson Laboratory as young adults and crossed in our animal facility. The *PV-Cre; Ai14* and *SST-Cre; Ai14* mice were heterozygous for both *Cre* and *Lox-Stop-Lox-tdTomato* alleles and were bred in-house by crossing female homozygous PV-Cre or SST-Cre mice with males homozygous for the *Ai14* reporter gene, respectively. Heterozygous floxed Gabrd mice (Lee and Maguire, 2013) were generated by Dr. Jamie Maguire’s group.^17^ A pair of breeders were kindly gifted to us to establish an in-house bred colony. Services from Transnetyx were used for genotyping mouse lines.

### In-situ Hybridization Assay

Manual RNAScope assay was performed using the Multiplex Fluorescent Reagent Kit v2-Mm (ACDbio, 323136) with the RNAscope® Probe-Mm-Gabrd-C3 (#459481) and RNAscope® 2.5 LS Probe-Mm-Gad1-C2 (#400958). Mice were perfused with PBS and 4% PFA. Brains were post-fixed in 4% PFA overnight at 4°C and then dehydrated in 30% sucrose for a day. 15 µm sections were obtained using a cryostat and kept at −80°C before use. The staining procedures followed the manufacturer’s protocols for fixed-frozen tissue samples with minor modification. The tissue was digested in protease III for 10 min instead of 15-30 min.

### Immunohistochemistry

Aged-matched adult (*PV-Cre;Ai14 and* SST-Cre;Ai14) mice were anesthetized and transcardially perfused with phosphate-buffered saline (PBS), followed by 4% paraformaldehyde (PFA). Brains were dissected out and post-fixed in 4% PFA overnight at 4°C. The following day, brains were transferred into PBS and stored at 4°C for up to a week. Coronal sections (50 μm) containing GC were prepared on the vibratome (VT1000, Leica) and stored in PBS at 4°C for up to a week until staining. Slices from PV-Cre and SST-Cre lines were processed in parallel from perfusion through immunostaining, imaging, and analysis. For immunostaining, brain slices were washed in PBS 3 x 5 min at RT and incubated in 5% Normal Goat Serum (NGS) and 5% Bovine Serum Albumin (BSA) in PBS with 0.1% Triton-X-100 at 4°C for 1 hr to block excess protein sites on brain tissue. Following blocking, tissue was incubated overnight at 4°C in 5% NGS, 1% BSA in PBST (0.1% Triton-X-100) solution containing the primary antibody against δ-GABA_A_R (1:250, Alomone Labs, AGA-014). Slices were then incubated with Alexa-Fluor 488 conjugated goat anti-rabbit (1:300, ThermoFisher, A-11029) for 4 hr at 4°C followed by 30 min counterstain with Hoechst 33342 (1:5000, Invitrogen, H3570). Following 3 x 10 min washes in PBS, sections were mounted onto glass slides with Fluoromount-G mounting media (VWR, 100241-874). Immunolabeling was visualized on a Zeiss confocal and quantified using ImageJ and Cell Profiler software. Images were acquired in 1 μm steps using a 20x objective, and the laser power and camera settings were minimally adjusted to ensure a high signal-to-noise ratio for each sample. The image step with the highest immunoreactivity in the PV/SST channel (594 nm) was used to locate PV/SST positive cells and to verify that the δ colocalization measured in the respective channel corresponds to that same cell.

### Electrophysiology

Mice (2-5 months) were anesthetized with isoflurane using the bell jar method and rapidly decapitated. The brain was dissected in ice-cold, oxygenated standard Artificial Cerebro-Spinal Fluid (ACSF) and oxygenation was maintained throughout dissection and slicing procedures by bubbling ACSF with carbogen (95% oxygen and 5% carbon dioxide). Coronal slices (350 μm) containing GC were prepared using a fresh tissue vibratome (Leica VT1000), allowed to recover in 34°C ACSF for 30 min, and then brought to room temperature (RT) for 1 hr before beginning experiments. Individual slices were transferred to the recording chamber and perfused with an oxygenated ACSF. Whole-cell patch-clamp recordings were obtained from visually identified pyramidal neurons under differential interference contrast (DIC) optics using borosilicate glass pipettes with resistance of 4 to 5 MΩ filled with a cesium sulfate–based internal solution (see *Solutions* section below). Spontaneous inhibitory postsynaptic currents (sIPSCs) were recorded in voltage clamp by holding neurons at +10 mV, the expected reversal potential for AMPA and NMDA receptors-mediated currents. Cells with a change in series resistance >20% during recording were excluded from the analysis. Immunohistochemistry was used to confirm neuron identity *post hoc*.

### Post hoc labeling of recorded neurons

Slices from electrophysiological recording were post-fixed in 4% PFA after recordings for later confirmation of cell type. They were washed 3 x for 10 min in PBS at RT and then blocked in 5% NGS and 5% BSA in PBS with 0.1% Triton X-100 for 2 hr at RT. Slices were then incubated with the primary antibodies streptavidin Alexa Fluor 568 conjugate (1:2000; Invitrogen, S11226) and mouse anti-GAD67 (1:500; MilliporeSigma, MAB5406, monoclonal) overnight at 4°C in a solution containing 1% NGS and 1% BSA in PBS with 0.1% Triton X-100 and counterstained with Hoechst 33342 (1:5000, Invitrogen, H3570). Following 3 x 10 min washes in PBS, sections were mounted onto glass slides with Fluoromount-G. After drying overnight, sections were imaged with a laser scanning confocal microscope (Zeiss) at 10x and/or 20x magnification to validate cell type and location within GC.

### Solutions

ACSF contained the following (mM): 126 NaCl, 3 KCl, 25 NaHCO3, 1 NaHPO4, 2 MgSO4, 2 CaCl2, 14 dextrose with an osmolarity of 315-320 mOsm. The internal solution to assess spontaneous inhibitory events contained (mM): 20 KCl, 100 Cs-sulfate, 10 K-HEPES, 4 Mg-ATP, 0.3 Na-GTP, 10 Na-phosphocreatine, 3 QX-314 (Tocris Bioscience), 0.2% biocytin (reversal potential *(V*rev)[Cations] = 10mV). The pH was adjusted to 7.35 with KOH, and the osmolarity was adjusted to 295 mOsm with sucrose. 500 nM Allopregnanolone (Steraloids, QX205296) and 20 µM picrotoxin (Tocris Bioscience) were applied sequentially to activate the tonic current and assess the involvement of GABA_A_ receptors.

### Implants of infusion cannulae and virus injections

Mice aged 2-5 months were anesthetized with an intraperitoneal injection of dexmedetomidine (1 mg/kg) and ketamine (70 mg/kg). The lack of toe pinching reflex ensured a surgical plane of anesthesia. Mice were placed in a stereotaxic frame (Kopf) and positioned on a heating pad (DC temperature control system, FHC, Bowdoin, ME) to maintain the body temperature at 35°C. The fur on the head was shaved and bupivacaine (2.5 mg/kg) was administered under the scalp for local anesthesia following scalp disinfection with three alternating applications of iodine and ethanol. Ophthalmic ointment was placed on the eyes to prevent dryness. Small craniotomies were made over the left and right GC (AP: 1.2 mm, ML: 3.2-3.5 mm relative to bregma). Stainless steel guide cannulae (26 gauge, 5.0 mm, Plastics One) were lowered 1.7 mm vertically from the cortical surface and fixed to the skull with acrylic dental cement (Stoelting). A metal head post was implanted posterior to the cannulae for head restraint during infusion. Once the dental cement was completely dry, dummy cannulae were inserted into each guide cannula, and animals were allowed to recover for at least one week before the behavior experiments. After the surgery, Antisedan (atipamezole hydrochloride, 1 mg/kg) and lactated ringer solution (1 ml) were administered subcutaneously to reverse anesthesia and for hydration, respectively. Mice were given carprofen (5 mg/kg) daily for at least 3 days after surgery.

In experiments designed to test the involvement of δ-GABA_A_Rs in the modulation of sucrose preference, viral injections were performed before the cannula implant. A pulled glass pipette front-loaded with one of 3 viral constructs (AAV9-GFP-Cre, titer 1 × 10^13^ genome copies (gc)/ml; AAV9-CamkII-eGFP, titer 2.8 × 10^13^ genome copies (gc)/ml; AAV9-mDLXp-mCherry-T2A-iCre, titer 9.93 × 10^12^ genome copies (gc)/ml (Vector Biolabs)) was lowered into GC and a microinjection pump (WPI UMP3T-1) injected 300 nL of virus at 1 nL/s. Two injections (150 nL each) were performed at two different ventral-dorsal locations ((1.75-1.9 mm below the dura). After each injection, the pipette was left in place for 8-10 minutes before being slowly retracted. Mice were given carprofen (5 mg/kg) daily for at least 3 days after surgery and allowed to recover for at least 2 weeks before beginning water restriction and behavioral studies.

### Validation of cannulae placement or virus injection sites

Mice were anesthetized and transcardially perfused with PBS, followed by perfusion with 4% PFA in PBS. Brains were collected and post-fixed in 4% PFA at 4°C overnight following post-fixation, then rinsed 3 x 5 min with PBS. Thin, 50μm coronal brain slices containing GC were cut on a vibratome (VT1000, Leica), washed 3 x 5 min in PBS at RT and then incubated with Hoechst 33342 (1:5000, Invitrogen, H3570) in PBS for 30 min at RT. Brain slices were rewashed with PBS before mounting onto glass slides with Fluoromount-G. Slices containing cannulae tracks and/or virus signals were mounted and imaged on a fluorescence microscope (Zeiss) at 10x magnification.

### Intra-GC bilateral Infusion

Mice in all behavior paradigms follow the same infusion protocol. On both control and drug days, mice were simultaneously infused with 1.6 µl of solution in each hemisphere using an infusion pump (Harvard apparatus 11 Plus 70-2212) at a 0.8 µl/min infusion rate. On control day, mice received an injection of 0.0007% DMSO (Tocris, catalog #3176) in saline, whereas on drug day they received a 500 nM allopregnanolone (ALLO) dissolved in saline.

### Taste stimuli

Sucrose solutions were prepared daily using reagent-grade chemicals (Sigma-Aldrich, #S0389) dissolved in deionized water and presented at room temperature.

### Brief access test (BAT)

A Davis rig gustometer (Med Associates Inc., Davis Rig for Mouse-16 Bottle) was used to monitor licking behavior and to obtain sucrose preference curves. The device consisted of a chamber (14.5 cm wide, 30 cm deep, and 15 cm tall), a motorized moving table to deliver multiple solutions, a motorized shutter door to allow limited access to one tastant at a time, and computer and accompanying software to control tastant delivery and to record the timing of licks. The licking behavior was measured by recording the number of licks, the latency to the first lick, and inter lick intervals (ILIs) for each trial. For analysis, ILIs < 60 ms were removed to avoid inflation of lick counts by double contacts ensuing from a single tongue protrusion as adapted from Glendinning *et al*.^18^ An analog-to-digital converter and computer system registered licks by recording the change in capacitance induced by the tongue’s contact with the sipper tube. The apparatus was cleaned with 70% ethanol and rinsed with distilled water between mice.

Water bottles were removed from the home cage a day before training began. Each training session was 30 min, followed by free access to 1.5 - 2 ml of water. Mice were weighed daily to ensure they maintained at least 80% of their initial body weight. On day 1, mice were habituated to the Davis rig gustometer in a session where the shutter door remained open and allowed free access to one water bottle. Spout training with ad libitum access to one water bottle continued on day 2.

On day 3, mice were trained to the opening and closing of the shutter door (shutter training), which mimics the subsequent sessions’ trial structure. This session consisted of trials in which the shutter door opened, granting access to a bottle containing water. The shutter door closed after 15 s if no licks were detected, followed by a 5-s intertrial interval (ITI). If a lick was detected, the shutter door remained open for 10 s following the first lick. At the end of the 10 s window, the shutter door closed. The motorized table moved to present a different water bottle during the 5 s ITI. On days 4, 5, and 6, mice were trained to acquire sucrose preference curves in the gustometer. Days 5 and 6 began with mice being acclimated to head fixation and insertion of internal for infusions. On day 7, mice received a GC infusion of saline and on day 8 an infusion of allopregnanolone (ALLO). At the end of day 8, mice were returned to ad libitum access to water in their home cage, concluding the experiment. To obtain sucrose preference curves, 4 bottles containing different concentrations of sucrose, and 1 bottle of water were presented in a pseudorandom order. The range of concentrations was presented as a block, and the software randomized the presentation (without replacement) within each block. Sucrose concentrations (in mM): 10, 50, 200, and 600.

### Data analysis and statistics

We chose appropriate parametric or non-parametric statistical tests and accompanying post hoc analysis following the normality test of the data sets. Details about tests applied to data are indicated in the corresponding Results section.

### Image analysis

Images were segmented in ImageJ and analyzed using CellProfiler (version 4.2.6), an open-source software that uses advanced statistical algorithms to analyze biological images quantitatively. PV/SST cell bodies within each image were identified as objects and related to secondary objects, delta puncta. The puncta number per cell was collected per slice and averaged (3 slices per mouse). The pipeline consists of a background subtraction module to reduce noise, identifying primary objects (nuclei, cell bodies, and d subunit) and relating the objects to extract cell type-specific colocalization and overlap coefficient measurements. The highest PV/SST immunoreactivity cells were used to determine the co-expression of δ-GABA_A_R puncta. The overlap coefficient, Manders coefficient using Coste’s auto threshold, was used to assess the significance of the co-expression.

### Electrophysiology Analysis

Postsynaptic currents and synaptic events were detected and quantified using Easy Electrophysiology v.2.7.1 (Easy Electrophysiology Ltd), and the statistical analyses were done using GraphPad Prism 10 v. 10.2.3. Shifts in the holding current were calculated by taking current averages in 10 ms windows at baseline, in the presence of allopregnanolone (ALLO) and the presence of picrotoxin (Ptx). These windows were positioned within 90 s of the onset of the ALLO application, but the windows were adjusted to avoid overlap with noise or artifacts. Current deflections induced by the drugs were visually identified and confirmed with all points histogram of 60 s traces within each condition. To measure the kinetics of current events, current traces were first filtered with a 1 kHz eight-pole Bessel filter to denoise and visualize synaptic events. An event template was generated and used to pick events; the amplitude, rise, decay, and frequency of the first 100 events in each condition were averaged and plotted.

### Behavior Analysis

BAT data were analyzed using GraphPad Prism 10, version 10.2.3. The tastants/water (T/W) lick ratio comprised the number of licks in each trial divided by the average licking for each group’s (control and drug) water (0 mM sucrose) trials. This helps account for individual differences in licking behavior unrelated to the treatment but to inherent variability among subjects. A T/W lick ratio of 1.0 indicated similar licking for water and sucrose, thus no preference for sucrose. A two-way ANOVA was used to compare the sucrose concentration and drug effects. A sigmoidal 3-parameter logistic function [Y = b+((a-b) / 1+10^(logEC50-X)^] was applied to analyze differences between the saline and ALLO curves. Parameters in the equation: Y is the lick response, b is the minimum asymptote of the curve, a is the maximum asymptote of the curve, EC50 is the concentration of the agonist that produces a response halfway between the top and bottom and X is the logarithm of the concentration of the agonist.

